# CytoPro: A computational platform for accurate and robust assessment of cell contributions in bulk expression from diverse tissues and conditions

**DOI:** 10.1101/2021.08.19.456930

**Authors:** Adir Katz, Renaud Gaujoux, Hadas Orly, Elina Starosvetsky, Roye Rozov, Yoni Rabinovitch, Benny Halberstadt, Liron Bar, Yael Spector, Naama Wald, Chanan Rubin, Yair Benita, Shiran Gerassy-Vainberg, Shai S. Shen-Orr

## Abstract

Cells are the quanta unit of biology and their relative composition in a tissue is the prime driver of bulk tissue gene expression variation. When there is no cell information, deconvolution is an effective tool to achieve cell resolution, which provides important information for learning disease complexity and its interactions with treatments, drugs and/or the environment in a wide variety of contexts. Here we present CytoPro, a production-level tissue and condition-specific deconvolution platform, based on a large collection of human tissue-specific signatures derived from single and sorted cells. CytoPro infer per-sample multiple cell-type composition, given input bulk gene expression. CytoPro includes a rigorous QC pipeline for learning, generating and selecting signatures and performs internal automated validation using multiple QC test criteria including: Comparison to ground truth cytometry and pure sorted cells data, performance evaluation using simulated data including robustness to noise as well as agreement with biological expectations in validation datasets regarding genes and cells. We demonstrate that CytoPro outperforms existing deconvolution tools, in both accuracy and robustness. By exploring multiple datasets with predefined disease phenotypes, and analyzing a use-case of biological treatment response, we show the ability of CytoPro to flush out relevant cell biology in real pathological conditions.

## Introduction

Cells are major drivers of gene expression, and a key component for learning disease complexity and its interactions with treatments, drugs and/or the environment. Analyzing bulk gene expression, without accounting for variation in cell-type composition, may lead to considerable confounding effect. Current methods to obtain cell information include standard physical sorting methods as FACS, cytometry by time of flight (CyTOF), as well as single cell RNA sequencing (scRNA-seq). However, cytometry may suffer from experimental artifacts related to sample processing, doublets, gating and poor markers; whereas scRNA-seq has its own issues of high costs for studies with large sample size, sparsity, high sampling variation and difficulties to detect specific cell types such as neutrophils, neurons and adipocytes, whose dissociation by standard protocols may disrupt their cytoplasmatic content^1–3^. Ultimately, more often than not and especially in clinical trials where logistic burden is high, cell level data is not collected on subjects, yet samples for bulk whole genome gene expression are collected and measured.

Computational cell deconvolution methods allow inference of cell subset composition in a sample with the starting input being of a large set of genes (mRNA expression) measured in bulk tissue. Deconvolution methods can broadly be divided into enrichment-based and model-based methods, such as xCell^4^ and CIBERSORT^5^ respectively, among others. Although successfully employed in several applications, enrichment-based methods may suffer from lack of independence, while model-based methods may encounter co-linearity problems. Many of the currently available tools rely on reference data of predefined cell-types. Expanding this to detect additional cell-types and subsets may not be an easy task to perform using these tools, due to a lack of relevant available reference data.

Furthermore, in most cases, these signatures are generic and usually lack the biological context of the sample source or condition. This may impair the cell estimate accuracy and introduce bias, since the tissue specific environment and microenvironment as well as the biological condition may dictate cell specific expression^6–8^. Recently methods that rely on the availability of tissue specific relevant scRNA-seq reference data such as CIBERSORTx^9^ and Bisque^10^. Yet, these usually rely on methods a single or few datasets, whereas prediction based on multiple datasets from different experiments and platforms, may be more generalized and better representative of the population, thereby improving accuracy.

To overcome some of these limitations, we devised CytoPro, a new deconvolution platform that provides cell contribution estimates to evaluate per-sample cell content from bulk gene expression, that is tailored for tissue and condition specific settings. CytoPro includes a large collection of signatures generated from a large compendium of samples from multiple datasets of sorted cells and scRNA-seq. CytoPro has a large coverage of cell-types in total, where each signature includes only relevant cell-types, to improve performance and reduce false positive signal. An internal rigorous QC testing procedure automatically selects, generates and validates each signature according to multiple test criteria. We show that CytoPro outperforms existing refence methods in several performance tests, across different tissue sources and conditions. This platform may serve as a useful analytical framework in pharmaceutical development projects across multiple disease areas and diverse disease-relevant tissues.

## Results

### Method overview and downstream usage

Using bulk gene expression data as input, CytoPro provides sample-level estimation of multiple cell type contributions by using tissue and/or condition specific settings. We devised the CytoPro platform by learning, generating and selecting tissue-specific signatures, which are used by a deconvolution method to quantitate per-sample cell content (Fig 1a). To generate and select a best performing tissue- and cell-type-specific signature, we followed an internal multi step process: 1) selection of appropriate cell-types per tissue and identification of relevant datasets from a large manually curated compendium of 9000 samples from RNA-seq and expression arrays comprised of sorted cells from different tissues and conditions and single cell datasets collected from peer-reviewed publications and proprietary sources. 2) signature generation and selection based on machine learning framework which produces hundreds of candidate signatures, and selection of the best performing one using training, test and validation sets. 3) SiQC (CytoPro Signature Quality Control) which performs automated signature validation using multiple QC test criteria. QC criteria include: (a) correlation with ground truth cytometry proportions across different datasets, (b) cell type specific signature overlap with literature curated markers, (c) gene expression-cell estimates correlation assessment, (d) performance in samples of sorted cells tested against multiple held-out datasets, (e) performance evaluation by *in-silico* simulations of complex mixtures generated from sorted cells, and (f) comparison to disease expectation using a pre-defined contrast, by direction and significance across different disease specific datasets. This QC process allows for high performance verification of the deconvolution method in both the statistical and biological aspects.

**Figure 1.**
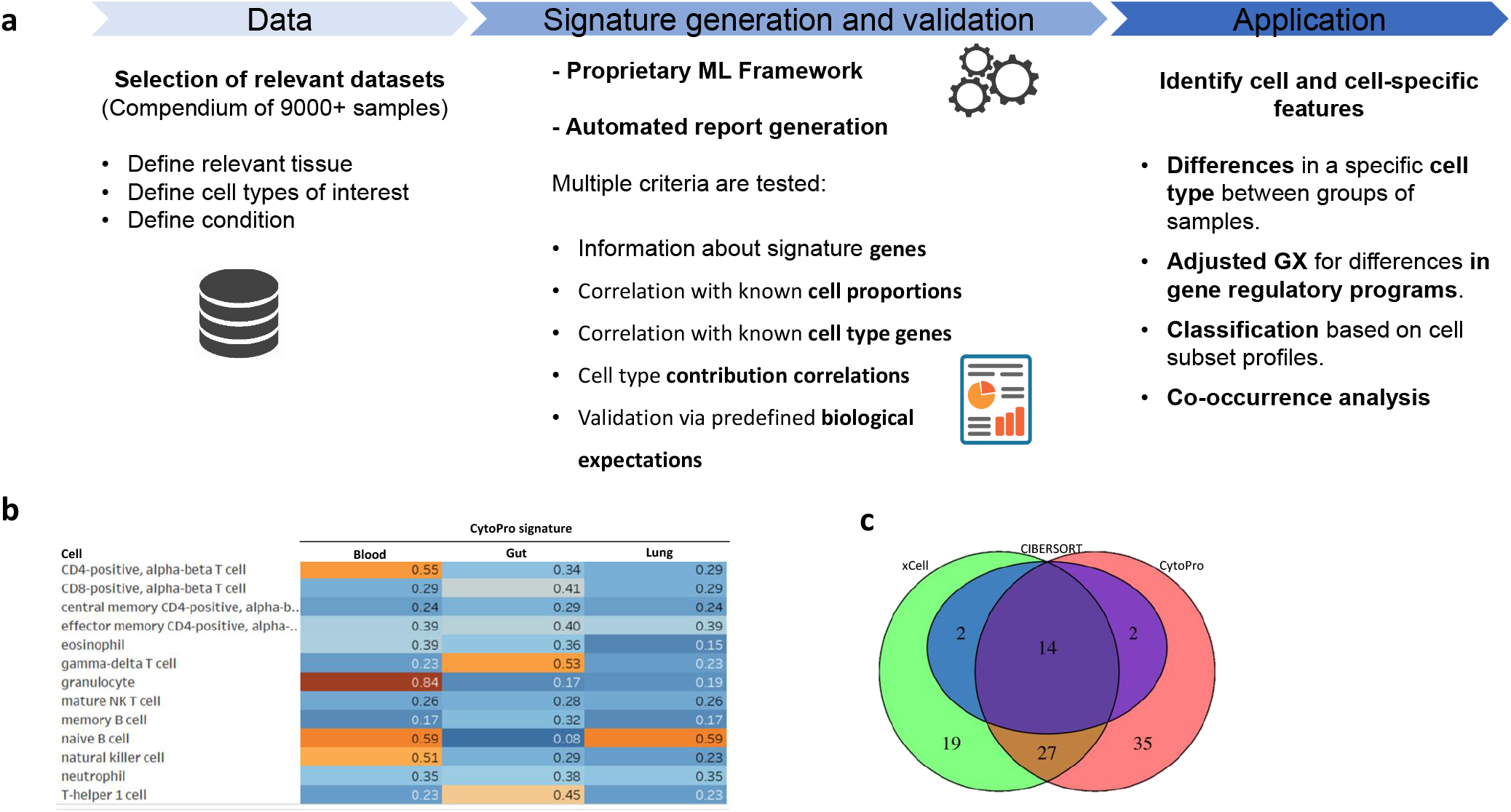
CytoPro workflow, coverage and application. **a**, CytoPro platform development and application involves several steps including data selection from a collection of datasets according to relevant tissue and cell-types. The signature generation process is based on proprietary machine learning framework which produces candidate signatures and selects the best performing one. This is followed by QC process to validate the selected signature by multiple test criteria, both by itself, and compared to reference methods. The QC results are summarized by SiQC report. After approval, the signature can be applied to a wide variety of downstream analysis. **b**, Heatmap summarizing the distinction rate of the CytoPro signatures from different tissue sources including blood, gut and lung. **c**, Venn diagram showing coverage of cell types by CytoPro and the two reference methods, CIBERSORT and xCell. CytoPro cell-state subsets by single-cell-based signatures and tumor signatures were not included in the analysis.

As immune cells located at different tissues, may have different expression profiles that are shaped by the tissue microenvironment, it is of high importance to generate tissue-specific rather than general signatures to avoid tissue bias and compromised cell estimate accuracy. This issue was addressed recently for mouse samples^6^ but not yet reported in humans systematically using a large number of tissue-specific datasets. Currently, the CytoPro platform contains a collection of tissue-specific validated signatures from different sources including blood and PBMCs (peripheral blood mononuclear cells), gut, synovium, lung, liver, myocardium and skin. In addition, CytoPro includes signature collections for tumor tissues comprising of kidney tumor signature designed for renal clear cell carcinoma, including primary tumors and metastases; bile duct tumor; liver tumor signature designed for hepatocellular carcinoma and angiosarcoma liver tumors; skin tumor signature designed for skin tumors including basal cell carcinoma, squamous, cell carcinoma and melanoma; several epithelial tumor signatures designed for general epithelial tumors which are not covered by tissue specific epithelial signature, epithelial tumor myeloid T-cells signature which provides high resolution cell contribution information of T cells and myeloid cells infiltrates, and epithelial ovarian cancer; and finally a pan tumor signature designed for solid tumors of various sources, including primary tumors and metastases. CytoPro signature collections cover a total number of 243 cell types, however for comparative purposes, following exclusion of single cell based or tumor signatures, we restrict ourselves in this work to 78 cells comparable with either CIBERSORT or xCell (Fig 1b-c and Table 1 for cell type list).

**Table 1.**
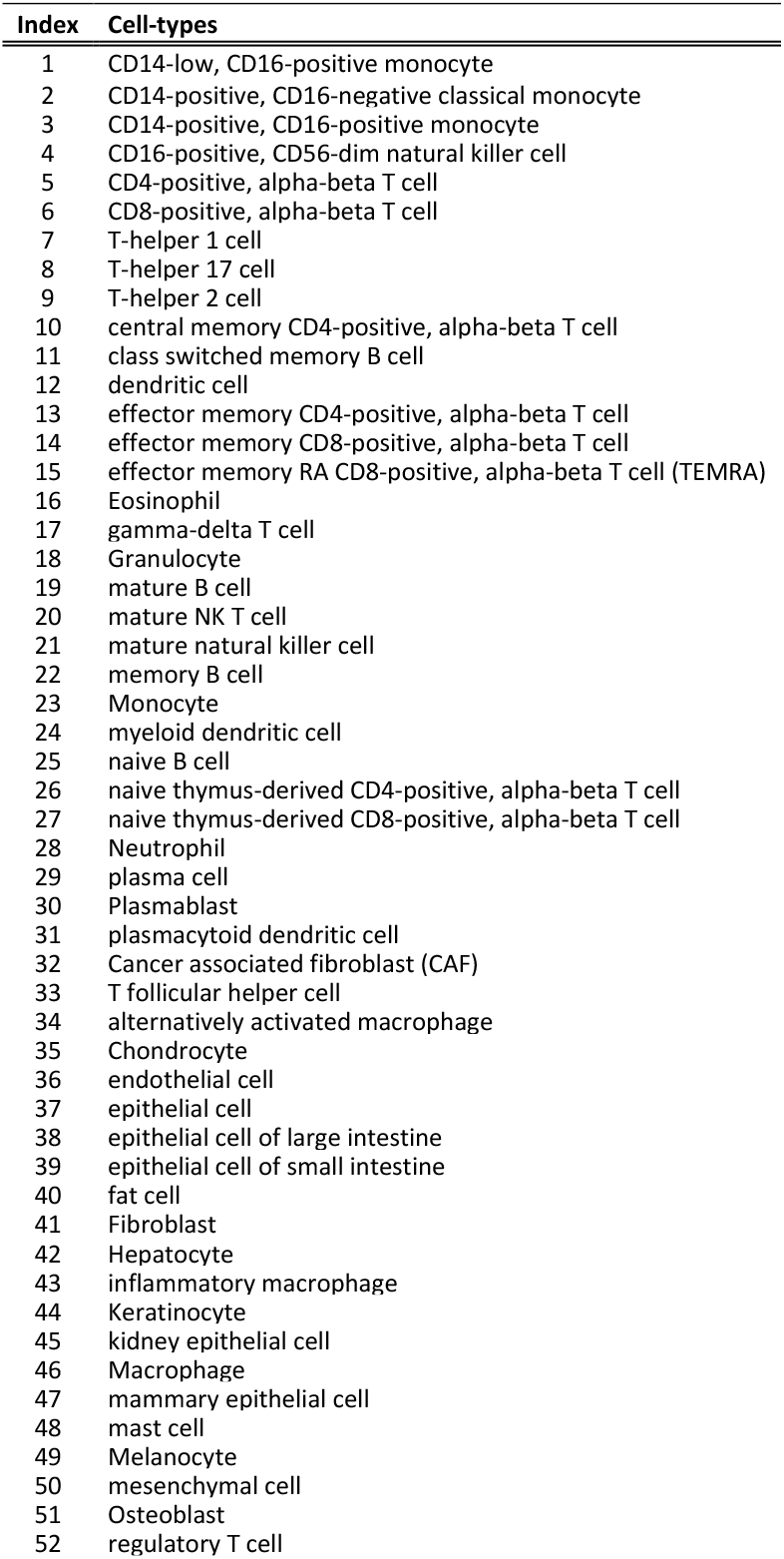
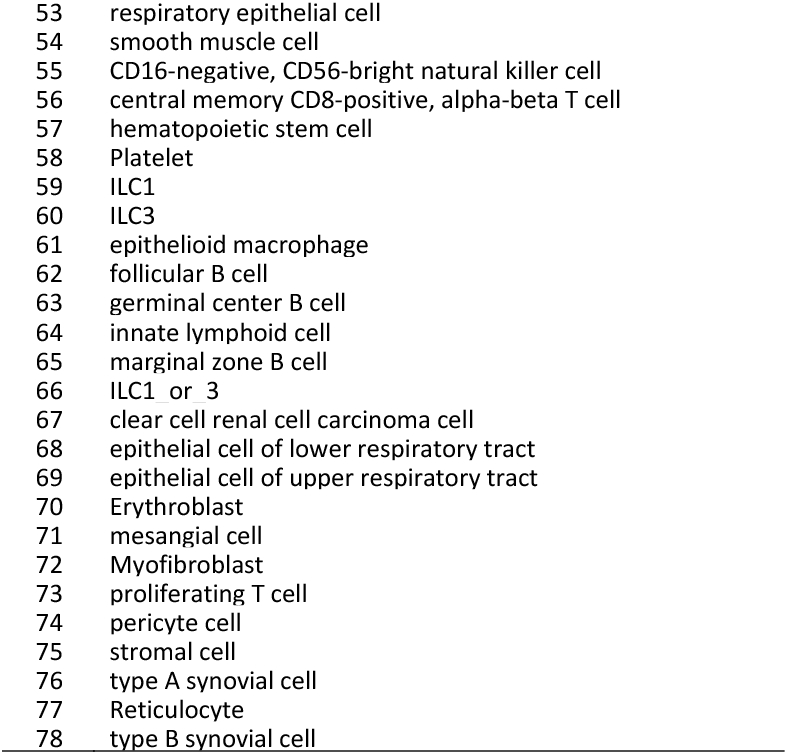
List of unique cell types covered by CytoPro, excluding single cell-based and tumor signatures.

The number of genes in a specific signature per cell type may range from dozens to hundreds, depending on the signature predictive model. To avoid high risk of false positive signal, CytoPro generates signatures only for relevant cell types in each specific tissue. Consistent with previous reports, our analysis confirmed the distinctness of signature genes across tissues per cell-type with a mean overlap of only 30% across distinct non-tumor tissues, indicating the importance of tissue-adapted functions. As representative tissues, we selected three different sources including blood, intestine and lung. Notably, the majority of cell-types presenting highly unique profiles were found in blood, likely due to the lack of non-immune cell subsets. We observed high distinction rate in granulocytes in blood compared to other tissues (84%) as well as naïve B cells (59%), CD4 positive, alpha-beta T cells (55%) and natural killer cells (51%) in blood (Fig 1b).

CytoPro outputs cell contribution scores which are a proxy for the cell type proportion in a sample. The score is a non-decreasing monotone value, which is correlated with the actual cell proportion and captures variation across samples, thereby suitable for direct identification of differences between groups of samples. CytoPro cell contribution scores can further be used for additional downstream analyses, giving users the opportunity to explore disease complexity and its interactions with any perturbation in a wide variety of contexts. Cell contribution scores may be used to perform cell-centered analysis to identify changes in transcriptional programs by accounting for the variation in cell composition when analyzing gene expression for variation in cell contribution scores. This procedure allows to place focus on detection of differences between conditions of the gene regulatory programs the cells are undergoing rather than those differences detected due to cell compositional differences, thereby unmasking additional signal^13^. Co-occurrence model can be applied to understand the underlying cell subsets network structure. This data can also be used for stratification or sample classification based on the cell-estimates profile (Fig 1a).

### Validation of CytoPro performance by direct comparison to ground truth cytometry datasets

To demonstrate CytoPro performance we compared estimated cell contributions to valid measures of cytometry proportions derived from blood. We tested nine relevant blood-based paired datasets of bulk gene expression from Affymetrix and Illumina platforms and cytometry data from both flow cytometry and CyTOF. Overall, we analyzed 674 samples. To objectively evaluate CytoPro performance, we tested its concordance with cytometry proportions against CIBERSORT^14^ and xCell^4^, two popular tools which represent model-based and enrichment-based methods respectively (Fig 2a). Only cell-types shared by both CytoPro and the additional reference method in each comparison were presented. We found that CytoPro compares well with the two methods on the tested datasets and shows the highest median correlation with cytometry proportions across the 14 shared canonical immune cell populations (Spearman’s r = 0.475, 0.359 and 0.364 for CytoPro, CIBERSORT and xCell respectively, n=100 bootstrap samples for determination of per-dataset and cell-type correlation, Fig 2b). Taken together, CytoPro contribution scores are overall correlated with actual cell type proportions. While they are not always strictly linearly associated with the actual cell proportions (Supp. Fig 1), they perform better compared to existing tools.

**Figure 2.**
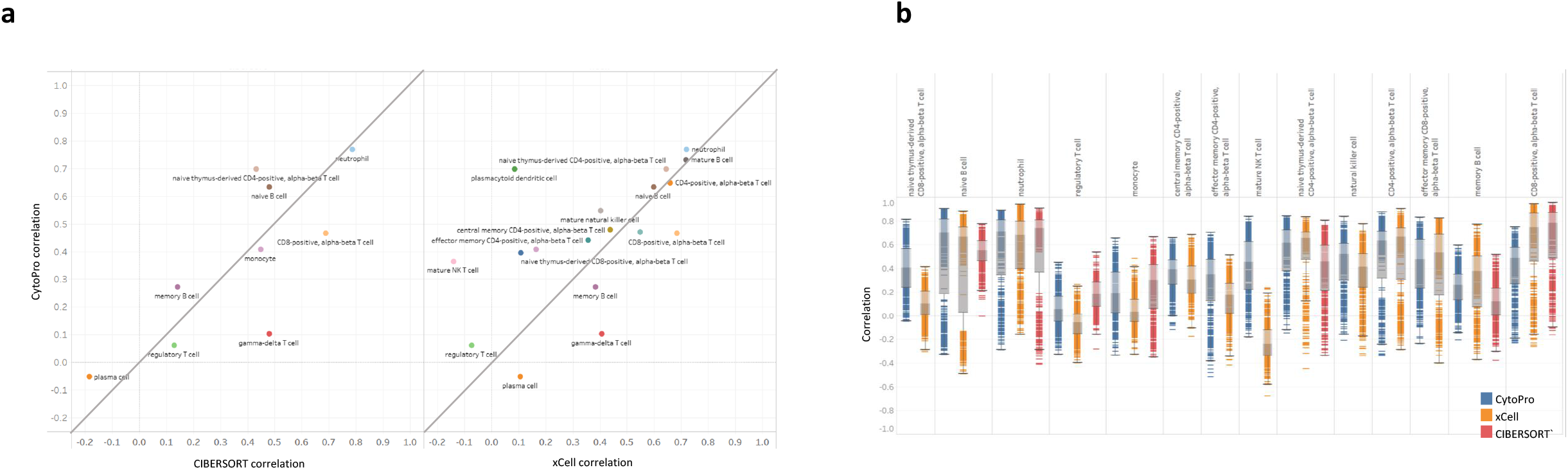
CytoPro performance by comparison to ground truth flow cytometry proportions in blood, across different datasets. **a**, Scatterplot presenting correlation of cell contributions of CytoPro to flow cytometry proportions and comparison to existing tools across 8 different paired datasets of cytometry and gene expression data. Spearman’s correlation coefficients with bootstrapping are shown (n=100). **b**, Boxplots of CytoPro correlations with cytometry measures per-cell type, compared to reference methods.

### CytoPro performance based on pure samples and *in-silico* simulated mixtures

As an additional confirmatory stage, and as a part of its signature selection process, we evaluate CytoPro cell-type specific signature performance in pure samples of sorted cells which are not part of the training process. We determine performance by quantifying the relative distance between the known true cell-type contribution score and the next cell highest score. A distance value of 1 indicates a perfect performance, while negative distance suggests incorrect identification where the highest contribution score is not assigned to the true cell type. To demonstrate performance according to this measurement, we evaluated the relative distance score of 37 different cell type in blood (Supp. Fig 2). All relative distances were positive, affirming correct identification of the true cell type compared to other cell types across different datasets.

CytoPro QC pipeline also includes performance tests on simulated bulk mixtures, generated from held out sorted cells expression data that were not used to train the classifier for the signatures generation. In this context, CytoPro evaluates performance in different conditions including variable number of cell types in the mixture, variable fractions of an unknown cell type that is not represented in the signature and sensitivity to input noise. For each cell-type-specific signature construction, we generated 100 synthetic mixtures. To do so, we sampled the proportions/weights of the tested cell-type from uniform distribution with a range of 0-1. The residual proportion was divided between the other cells in the mixture by sampling from dirichlet probability distribution, which has been commonly applied to model compositional data^15^. We created mixture bulk expression by multiplication of the weights with the centroid expression of the relevant sorted cell types. After applying the deconvolution algorithm to the synthetic bulk expression of the mixtures, we tested the concordance between the predicted cell contribution scores and the predetermined weights using Biweight midcorrelation (bicor) method with bootstrapping (n=500 bootstrap samples), which is a median-based correlation measurement that is robust to outliers^16^. We noticed only minimal change in the measured correlation as a result of changes in the number of cell-types in the mixture, with average absolute change of 0.01 and 0.03 in correlation at n=10 and n=20 relative to n=5 respectively (Fig 3a). Our results indicate that CytoPro performance remains stable and independent of the number of cell-types in the estimated sample.

**Figure 3.**
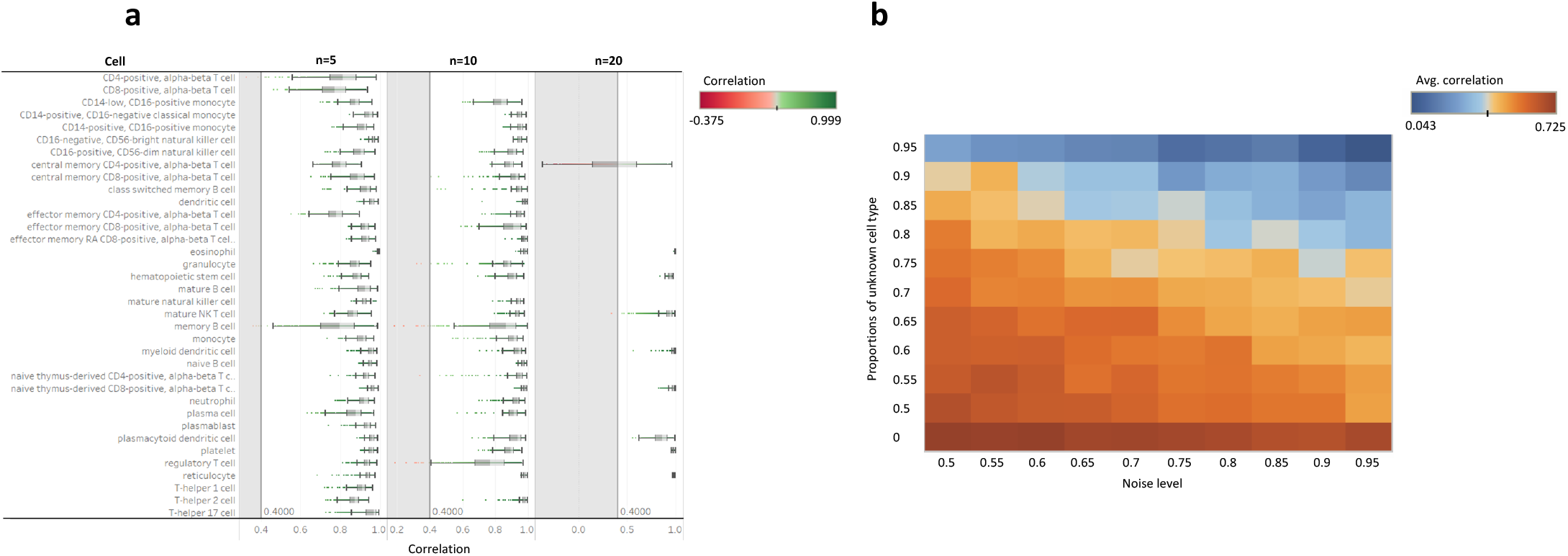
CytoPro robustness evaluation using simulated data. **a**, Evaluation of CytoPro performance under variable number of estimated cell types in a mixture (n=5, 10 or 20 different cell-types. Correlation values of the CytoPro cell-predicted estimates with the pre-determined weights per cell-type are shown. Values were calculated by the bicor method with bootstrapping (n=500). **b**, Heatmap representing the effects of a random spike-in of unknown cell-type (rows) and input expression noise (columns) on CytoPro performance, evaluated by the bicor measurement.

To assess the robustness of CytoPro performance to random noise that can stem from different experimental sources including experimental handling, measurements errors and differences between platforms^17^, we tested the performance of CytoPro on 100 simulated mixtures with increasing random noise. We introduce the random noise to the mixture bulk expression following sampling from normal distribution (∼N(0,1)) which was gradually increased through multiplying by an amplitude factor that ranged from 0 to 1, with sampling intervals of 0.2. By evaluating the correlation between the predicted and the predetermined weights along the magnitude of the tested random noise, we observed that CytoPro is robust and tolerates input variation and noise across cell-types (Fig 3b columns; for n=4 cell-types shared by CytoPro signatures and the reference existing tools, correlation by the bicor method).

On the same data, we also evaluated CytoPro performance in a simulated condition where an unknown cell-type which is not included in the tested signature, contributes to the mixture expression. We combined simulated bulk expression of known cell-types (as described above) with expression originating from increasing proportions of an unknown cell-type. We produced representative expression of the unknown cell-type by random selection of values from different genes, samples and cell-types of the reference sorted cells dataset. Based on the simulated data, CytoPro presents a stable performance along a variable range of proportions of an unknown cell population (Fig 3b rows; n=4 cell-types shared by CytoPro signatures and the reference existing tools, correlation by the bicor method). This further demonstrates CytoPro flexibility in tissues where not all cell-types are identified ahead.

We further assessed how CytoPro performs compared to existing tools based on the simulated mixtures data. To demonstrate its performance in additional tissue source, we focused on CytoPro lung signature. We found that CytoPro outperformed xCell for many cell types (Fig 4a, right) with the most prominent cells being T helper 1 cells, macrophages, mature NK T-cells and memory B cells, presenting large performance differences in favor of CytoPro (Fig 4b). CIBERSORT presented similar performance to CytoPro, yet it addresses fewer common cell-types using its standard LM22 signature (Fig 4a, left and Fig 4c). Evaluation of performance in simulated data compared to the reference tools, indicates that CytoPro keeps predicting cell contribution scores with high correlations to the predetermined weights, even when both variation sources of noise and the presence of an unknown cell-type are increased (Fig 4d, n=4 cell-types shared by CytoPro signatures and the reference existing tools, correlation by the bicor method). Taken together, the simulated data show that CytoPro is robust to potential biases and different sources of variations in the data including random noise, unknown cell population and varying number of estimated cell-types, demonstrating high performance compared to existing tools.

**Figure 4.**
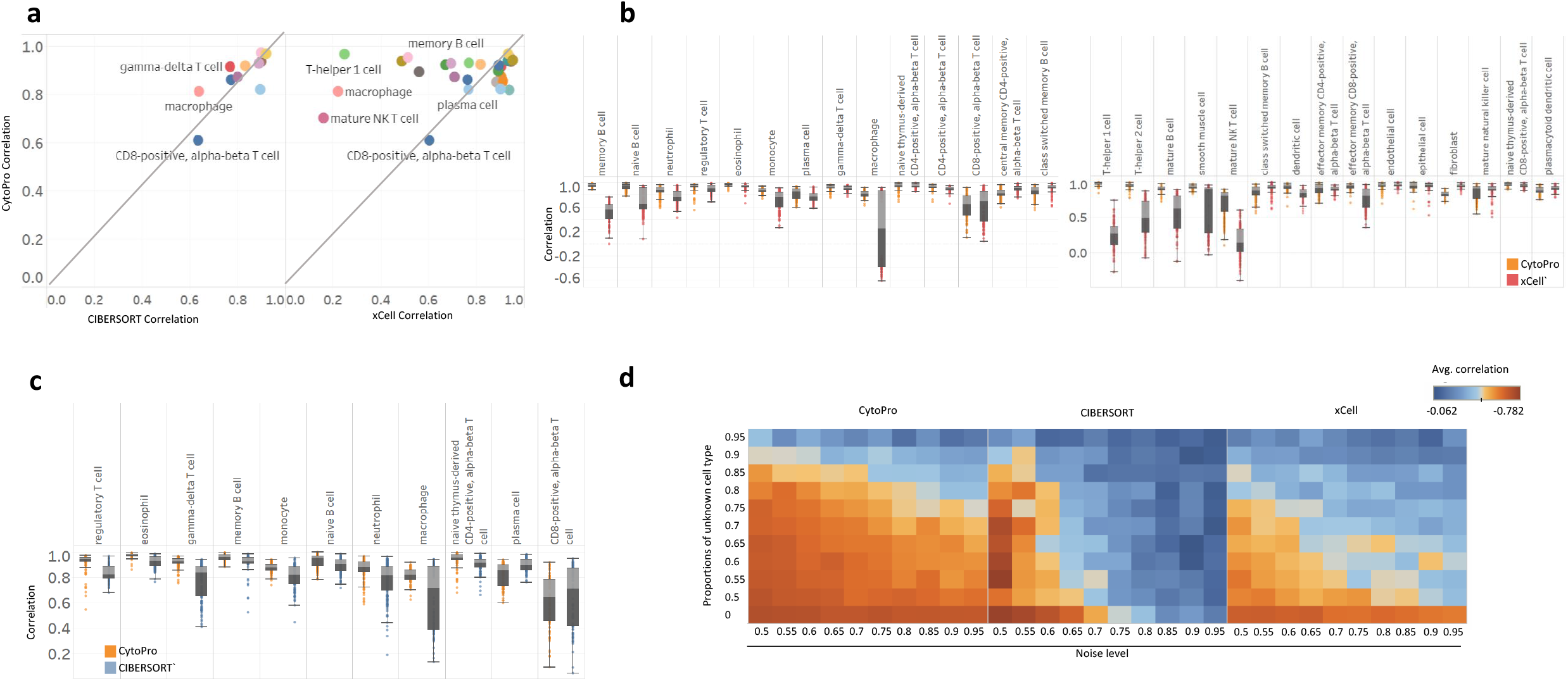
CytoPro performance comparison to existing tools using simulated mixtures. **a**, Scatterplots comparing CytoPro correlation with the simulated pre-defined weights using the CytoPro lung signature, to performance of the reference methods. Bicor values are shown. Only cell-types common to CytoPro and the compared reference method are presented. **b**, Boxplots of per-cell type CytoPro correlations over the simulated pre-defined weights, compared to the xCell reference method. **c**, Boxplots of CytoPro correlations with the simulated pre-defined weights, per-cell type, compared to the CIBERSORT reference method. **d**, Heatmap comparing the performance of CytoPro (left) and the reference methods, CIBERSORT (center) and xCell (left) under simulated input noise and varying fraction of unknown cell-type. Colors denote bicor measurement values.

### Validation of CytoPro performance by agreement with known biological expectations

An additional and essential stage in the QC workflow of CytoPro is comparison to known biological information to validate each modeled tissue-specific signature. In this context, CytoPro evaluates two main analytical aspects including comparison of cell-specific signature genes to known cell biomarkers and validation through knowledge-based expectations of held-out disease datasets using predefined contrast matrices.

The CytoPro platform includes a set of annotated cell-specific markers for each cell-type, curated from literature. We define both levels of mandatory markers by which cells can be uniquely isolated, and positive markers which are functional markers or markers presenting high data-driven correlations. CytoPro calculates Pearson correlations between relevant cell type contributions and the expression of the corresponding marker genes, for each dataset in the CytoPro validation-disease collection, and averages across all datasets. Accurate prediction of cell estimates based on valid signature is expected to yield stable positive correlations with the relevant gene set markers. Assessment of the correlation between representative cell types and their corresponding markers using the CytoPro gut signature in CD-related datasets as an example, confirmed positive correlation in almost all the tested markers, supporting known predefined biological expectations (Fig 5a). While xCell presents similar trends, CIBERSORT underperforms, by exhibiting negative correlations in several cell-types including macrophages, eosinophils and gamma-delta T cells.

**Figure 5.**
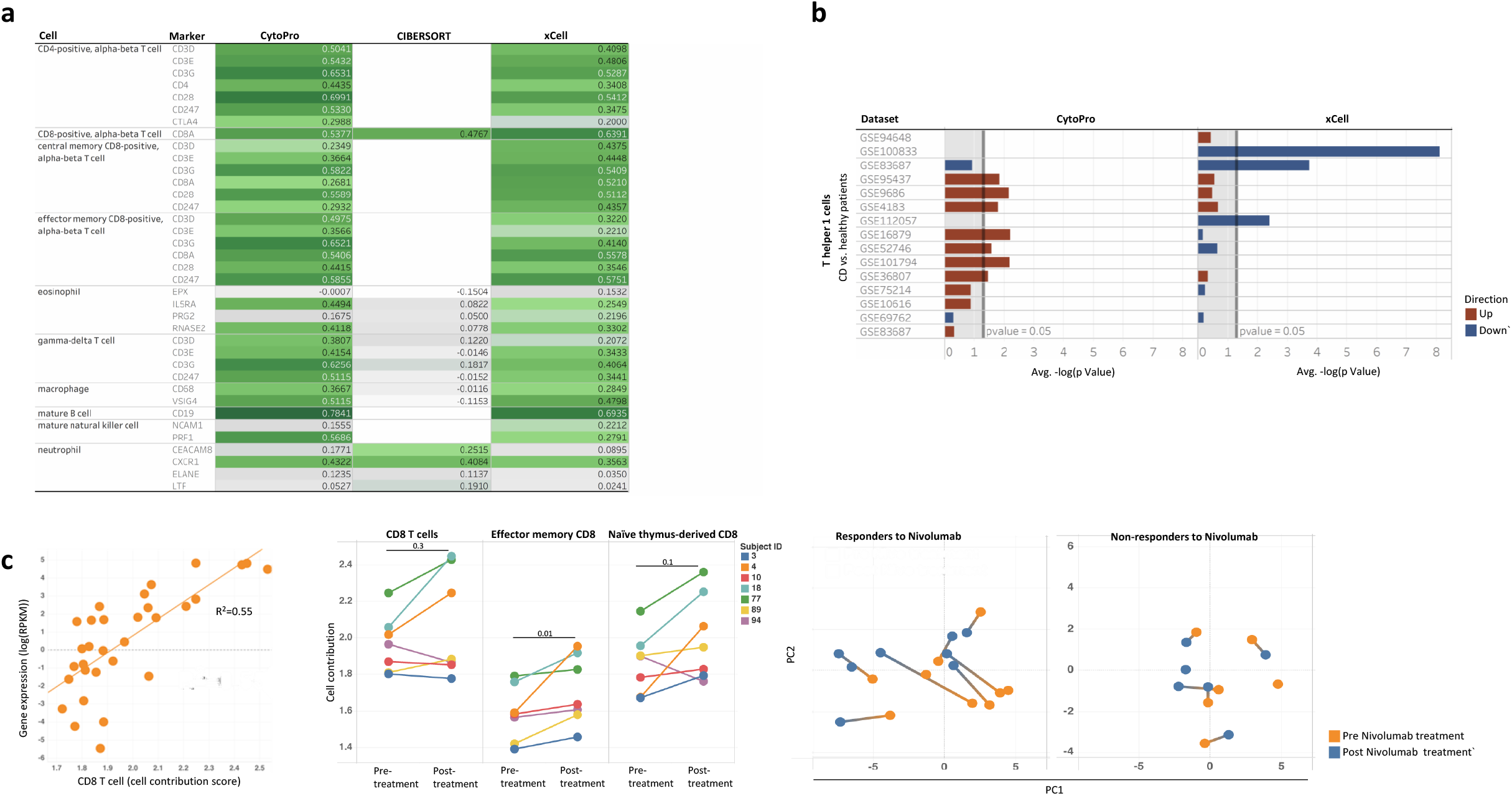
CytoPro evaluation by agreement with known biological expectations. **a**, Correlation of cell type contributions with known cell marker genes across representative cell-types for CytoPro gut signature and the existing reference methods in CD-related datasets from the CytoPro disease-validation collection. Average Pearson correlation coefficients across the relevant datasets are shown. **b**, Bar plots comparing predicted Th1 cell estimates differences between CD and healthy controls. **c**, CytoPro cell contributions detect expected treatment responsiveness.

To evaluate agreement with knowledge-based differences between disease states, CytoPro tests a pre-defined matrix of expected cell-types and tissue specific contrasts, related to diverse disease conditions including Crohn’s disease (CD), ulcerative colitis, rheumatoid arthritis, psoriasis, pancolitis, asthma, atopic dermatitis, idiopathic pulmonary fibrosis and NASH (Non Alcoholic Steato Hepatitis), which keep on expanding with additional cumulative data. Different disease states are tracked including disease vs. healthy control patients, paired diseased vs. healthy tissues (i.e. inflamed-non inflamed tissue, lesion – non lesion, tumor – adjacent uninvolved tissue etc.), active vs. remission, and treatment responders vs. non-responders. This contrasts matrix includes information on expected condition dependent-differences per-cell type, with a specified directionality. To measure performance according to this parameter, we compute a score based on the expected per-cell outcome and cell contribution given a tested signature, across all datasets. The computed score ranges between 0-1, where a value of 1 indicates that for the tested cell-type, CytoPro provided cell contribution scores that correctly identify expected differences across all the validation datasets by both significance and expected direction. As an example, we analyzed Th1 cells in CD, which were shown to accumulate in the intestinal tract of individuals with CD and are directly associated with disease^18^. Applying CytoPro platform using the gut signature, we successfully predicted increase in Th1 cells in intestinal tissue across multiple data sets (Fig 5b). Most differences were missed by xCell estimates, with some datasets even showing inversed differences compared to the expected direction.

### Application of CytoPro to real bulk expression data, recovered known biology

We also tested CytoPro cell content estimates in a real-life use case of pre- and post-anti-PD1 treatment (Nivolumab) in melanoma patients, using publicly available dataset (GSE91061^19^).

Expression of PD1 on CD8+ T cells was demonstrated to be associated with T cell exhaustion in melanoma, reflecting gradual loss of effector function during tumor progression^20,21^. Consistent with this, although PD1 is not a marker gene, it correlated with the CytoPro predicted CD8 cell contribution score in the tested samples (Fig 5c, left; R^2^=0.55). Furthermore, efficient treatment with anti-PD1 was shown to induce CD8 T cell proliferation^22^, leading to significant accumulation of CD8+ TILs (tumor infiltrating T cells) within the tumor. Comparing CytoPro contribution scores of different subsets of CD8 pre and-post treatment in samples of anti-PD1 responders revealed a significant increase in effector memory CD8 T cells post treatment (Fig 5c, center, p=0.01 by Wilcoxon test). This finding is corroborated by previous report showing that PD1+ effector memory CD8+ T cells controlled tumor growth and responded to PD1 blockade^23^. To further compare responders and non-responders treatment-induced cellular alterations, we looked at a contribution score-based PCA (Principal Components Analysis). Interestingly, we observed larger cellular differences in responders compared to non-responders (Fig 5c, right), indicating induction of immune response under efficient treatment of PD-1 blockade. These results signify the ability of CytoPro to identify relevant cellular compositional changes in real biological setups.

## Discussion

CytoPro provides per-sample cell contribution scores to evaluate cell composition based on human bulk gene expression in a wide variety of tissue sources and conditions. We demonstrated that CytoPro predicts accurate and robust cell estimates, via multiple performance tests including comparison to standard physical sorting methods as FACS and CyTOF, and comparison to sorted cell cultures; correlation of predicted estimates with predetermined weights using simulated mixtures, generated with different sources of variation including expression noise and varying proportions of unknown cell population; and also through comparison to expected biology including correlation with known cell markers and identification of expected disease related differences. We found that CytoPro consistently recovers expected trends of disease-associated phenotypes or treatment-expected responses, indicating that the cell contribution scores reflect relevant biological cell-derived processes and related clinical outcomes.

CytoPro compares favorably to existing tools which represent different reference-based deconvolution approaches (enrichment-based and model-based algorithms), in several evaluation criteria. The high performance of CytoPro is mainly attributed to three key elements. First, CytoPro deconvolution is tissue-specific and looks only for the relevant cell types in each specific tissue source. It currently contains multiple signatures designated specifically by tissue and tumor specific sources. Since cells have tissue-adapted functions and their expression profile is highly depended on the tissue-microenvironment, the development and usage of tissue-adapted signature for each relevant cell-type, has a decisive effect on the predicted cell contribution accuracy. Second, CytoPro has superior coverage of cell-types. CytoPro evaluates a large number of different cell-types at high cellular resolution. This enables to achieve a better performance and to decrease the probability for bias as a result of underestimating specific cell-types. Third, CytoPro platform is generated under a rigorous QC process which performs automated validation using multiple QC test criteria for each cell type in each signature, using a large collection of datasets for signature generation and validation.

CytoPro provides contribution score estimates for cell proportions, highly relevant for identification of biological differences between phenotypes or groups. The ability to infer correctly the actual true proportion of a cell, from deconvolution methods, popularly referred to as “absolute deconvolution”, has been raised in the academic community as achievable, however, it is our opinion that such approaches have limited accuracy, since any mistake in one cell-type propagates to others. In contrast, contribution scores are more reliable and can be trustily compared across samples. In addition, it requires a predefined signature and prior knowledge of the comprising cell-types in the estimated samples. Yet, considering the large collection of validated signatures in the CytoPro platform, which keeps growing and continuously being developed, it supports a wide range of tissue sources and biological conditions.

CytoPro cell estimates can be used to decouple two main dimensions of information from bulk gene expression data which makes the biological interpretation more focused and precise including differences in cell-type estimates, and differences in cell-regulatory programs by accounting for confounding effects of cell type proportions. Given its performance, robustness to different sources of variation, its ability to capture relevant biology and its high coverage of tissue sources, cell types and conditions, we believe that CytoPro platform constitutes a useful analytical framework to users for studying disease complexity and responses to drugs, treatments or environmental factors.

